# CRISPR spacers indicate preferential matching of specific virioplankton genes

**DOI:** 10.1101/487884

**Authors:** Daniel J. Nasko, Barbra D. Ferrell, Ryan M. Moore, Jaysheel D. Bhavsar, Shawn W. Polson, K. Eric Wommack

## Abstract

Viral infection exerts selection pressure on marine microbes as viral-induced cell lysis causes 20 to 50% of cell mortality resulting in fluxes of biomass into oceanic dissolved organic matter. Archaeal and bacterial populations can defend against viral infection using the CRISPR-Cas system which relies on specific matching between a spacer sequence and a viral gene. If a CRISPR spacer match to any gene within a viral genome is equally effective in preventing lysis, then no viral genes should be preferentially matched by CRISPR spacers. However, if there are differences in effectiveness then certain viral genes may demonstrate a greater frequency of CRISPR spacer matches. Indeed, homology search analyses of bacterioplankton CRISPR spacer sequences against virioplankton sequences revealed preferential matching of replication proteins, nucleic acid binding proteins, and viral structural proteins. Positive selection pressure for effective viral defense is one parsimonious explanation for these observations. CRISPR spacers from virioplankton metagenomes preferentially matched methyltransferase and phage integrase genes within virioplankton sequences. These viriolankton CRISPR spacers may assist infected host cells in defending against competing phage. Analyses also revealed that half of the spacer-matched viral genes were unknown and that some genes matched several spacers and some spacers matched multiple genes, a many-to-many relationship. Thus, CRISPR spacer matching may be an evolutionary algorithm, agnostically identifying those genes under stringent selection pressure for sustaining viral infection and lysis. Investigating this subset of viral genes could reveal those genetic mechanisms essential to viral-host interactions and provide new technologies for optimizing CRISPR defense in beneficial microbes.

---

Between 20 and 50% of microbial mortality within marine systems results from viral infection and lysis. As a consequence, these processes are critical in driving carbon and nutrient cycles within the sea (1, 2). In response to the substantial pressure of viral predation, a number of sophisticated defense systems have evolved within cellular microbial hosts including: alteration of cell surface receptors, production of extracellular polysaccharides (3), restriction modification systems (4), and the clustered regularly interspaced short palindromic repeat (CRISPR) system. Of these systems, the CRISPR system is perhaps the most adaptable and specific, acting as an acquired immune system in Bacteria and Archaea against bacteriophage and archaeal viruses respectively, as well as other invading foreign DNA, such as plasmids (5). The adaptability of the CRISPR system for targeting specific DNA regions for nuclease digestion has been leveraged into a new and powerful approach for selective genome editing within complex plant and animal genomes (6).

The CRISPR locus is comprised of CRISPR-associated (*cas*) genes and one or more CRISPR sequence arrays consisting of a repeating pattern of different spacer sequences and the same hairpin repeat sequence. It is the spacers that enable the adaptable and gene-specific inactivating mechanism of the CRISPR system. Spacers are short segments (26-72 base pairs (7)) of sequence that are homologous to phage or plasmid DNA. Each spacer is flanked by comparably sized repeat sequences. The repeats form a hairpin secondary structure and are conserved among bacterial and archaeal species. The number of spacers in a CRISPR array varies from 2 to over 200 (7) and, interestingly, the position of a spacer in the array can provide an historical timeline of viral host encounters (5).

After transcription, Cas proteins cleave repeats from the array transcript creating small interfering CRISPR RNAs (crRNAs). The crRNAs are comprised of one spacer flanked on either side by half a repeat. If a spacer sequence within a crRNA matches a segment of an invading virus’ genome, then the small interfering crRNA will target the genomic DNA or RNA for destruction by the Cas proteins thus preventing viral replication and ultimately cell mortality (8). Assuming that every gene a virus carries in its genome is essential for successful infection and lysis, then, successful CRISPR inactivation of any viral gene should prevent cell mortality from viral lysis. Given this understanding of CRISPR defense against viral infection, we should expect no preferential matches of viral genes to CRISPR spacer sequences. However, if there are differences in the effectiveness of inactivating certain viral genes over others, then certain viral genes may demonstrate a greater propensity to be matched by CRISPR spacers. This hypothesis was addressed by identifying spacers within microbial and viral metagenome sequence libraries and investigating whether subsets of viral genes were preferentially matched by these CRISPR spacers.

CRISPR spacers offer a powerful tool for investigating phage-host interactions as spacer sequences can link phage and host populations within complex microbial communities (9, 10). For example, this approach was used to identify the microbial hosts of unknown viral populations within the extreme environments of deep-sea hydrothermal vents (11, 12). The biochemical mechanism controlling the selection of protospacer sequences (i.e. candidate spacers from invading viral and plasmid DNA) relies on a short DNA motif (usually 2-6 base pairs) directly adjacent to protospacer sequences (protospacer adjacent motif or PAM) (13),(14). Because the PAM is a short sequence, these motifs can be common within a viral genome and thus, the PAM alone does not necessarily predispose particular viral genes as possible protospacer targets. However, positive selection for more effective viral resistance would mean that certain subsets of viral genes are preferentially represented as targets of CRISPR spacers within natural virioplankton communities. Information on viral genes preferentially matched by CRISPR spacers could indicate those viral genes most critical to successful viral replication and lysis. Given that the function of most viral genes is unknown (15), information on preferential spacer targeting could provide clues as to the subset of unknown viral genes that are under stringent selection for successful infection and host cell lysis. Fundamental information on the CRISPR susceptibility of particular viral genes could be leveraged to engineer more effective phage resistance in beneficial microbes.

Spacers can be identified within DNA sequence libraries based on their characteristic repeat-spacer pattern within a CRISPR array. Several tools are currently available for identifying CRISPR spacer arrays, however, these tools tend to have a high false discovery rate of spacer sequences as repeat sequence arrays resembling CRISPR spacer arrays are common within microbial genomes (16). To address this shortcoming CASC (CASC Ain’t Simply CRT) was developed as a discovery tool capable of validating the accuracy of CRISPR spacer predictions. CASC employs a modified version of the CRISPR Recognition Tool (CRT) (16) to identify putative CRISPR arrays followed by novel heuristics (search for known repeats, spacer size distribution check) to examine and validate each putative CRISPR array. CASC is able to run in an exploratory (liberal) mode, as well as a stricter (conservative) mode in which identified arrays must contain known repeat sequences or Cas protein genes near the array.

After validation, CASC was used to identify CRISPR spacers within large collections of marine microbial metagenome sequence data from the Global Ocean Sampling (GOS) and *Tara* Oceans expeditions (17, 18). These spacers were then used to examine phage-host interactions throughout the global ocean and identify common genetic vulnerabilities among viral populations exploited by marine prokaryotes to defend against viral infection.

## RESULTS

### CASC validation with artificial data

Two artificial metagenomes were created to simulate Illumina reads and pyrosequencing reads. Both of these metagenomes were comprised of the same ten bacterial genomes: five genomes containing CRISPR arrays and five genomes without CRISPR arrays (see methods section).

The simulated Illumina sequence reads (150 bp, paired-end) were assembled with SPAdes (19) and produced ca. 1,800 contigs (mean length of 17,700 bp). Only one of the ten genomes (*C. trancomatis* F/SW5) was completely assembled into one contiguous sequence. Although the remaining genomes were fragmented into many contigs, the known CRISPR arrays were represented in the assembled dataset. The second artificial metagenome was composed of ca. 1 million pyrosequencing reads (450 bp) that were directly analyzed without assembly.

Each CRISPR algorithm evaluated (CASC, CRT, PILER-CR (20), and CRISPR Finder (21)) performed better in terms of sensitivity (ability to detect spacer loci) and precision (ability to detect only valid spacer loci) when searching for spacers within assembled contigs from Illumina sequence libraries as opposed to pyrosequencing reads (Tables S2 and S3). CASC’s validation steps, which remove potentially spurious CRISPR predictions, resulted in more accurate CRISPR spacer predictions (Illumina contigs precision = 1.0; pyrosequencing reads precision = 0.82) than all of the other tools that were evaluated.

### Spacer predictions in GOS metagenomes

The GOS reads dataset provided spacers from a broad geographic cross-section of bacterioplankton communities. Because the GOS sequence reads averaged 915 nucleotides in length it was possible to search for CRISPR arrays within unassembled reads. CASC (in liberal mode) was used to search for CRISPR spacers in all read sequences from GOS. CASC identified 12,606 CRISPR spacers (>99% did not match known spacers) contained in 2,686 arrays coming from 90% of all GOS sites (Additional file 1). The site with the most spacers (13% of all spacers observed within the entire GOS dataset) was GS033 (Punta Cormorant Lagoon, Floreana Island, Ecuador), which was the most heavily sequenced site. The number of spacers found was normalized by mega base pairs of reads sequenced at that site. Sites with the highest normalized spacer abundance were often lakes or lagoons (seven of the top ten), with most having more than two spacers per mega base pair of sequenced reads.

Nucleotide position histograms of the forward and reverse compliment direction of each CRISPR repeat sequence were used as a means of post hoc testing of CRISPR spacer arrays identified as “bona fide” and “non-bona fide” using CASC (liberal mode). Repeats within bona fide CRISPR spacer arrays showed distinct positional nucleotide signatures, whereas repeats within non-bona fide CRISPR array repeats showed no discernible signature as each position had an equal occurrence of each nucleotide (Fig. S2). The presence of a distinct positional nucleotide signature in the CASC bona fide repeats was indicative of a collection of true and functioning repeat sequences within the GOS data.

### Spacer predictions in *Tara* Oceans metagenomes

*Tara* Oceans assembled contigs contained more than twice as many spacers (29,879; 95% did not match known spacers) as the GOS reads (Additional file 2), likely due to the greater sequencing depth and number of samples in the *Tara* Oceans dataset. However, calculating the frequency of CRISPR spacers per mega base pair of sequence data was confounded by the fact that these data were collected from assembled contigs as opposed to single unassembled reads. To overcome this, read recruitment information was obtained for each *Tara* Oceans contig which enabled normalization of spacer abundance within the dataset (see methods). Between 15 and 71% of read bases were successfully recruited to contigs among the 178 *Tara* Oceans microbial metagenomes (Additional File 2). The fraction of each library associated with CRISPR spacers varied from 1 × 10^−4^ to 5 × 10^−8^ (Additional File 2).

After normalizing for sequencing effort, normalized spacer abundance (NSA) within the *Tara* Oceans metagenomes showed a positive correlation with sample depth (Pearson r = 0.42, p-value = 4e-9) (Fig. 1). The sample with the highest normalized spacer abundance was 122_MES_0.45-0.8, a mesopelagic sample having nearly 5,000 spacers per read Gbp recruited. Indeed, many of the samples with high NSA were from the mesopelagic zone (21 of the top 30). NSA showed a positive correlation with GC content as well (Pearson r = 0.51, p-value = 1e-13), which was not surprising to see as GC content also correlated strongly with depth (Pearson r = 0.74, p-value = 2e-16).

**Fig. 1.**
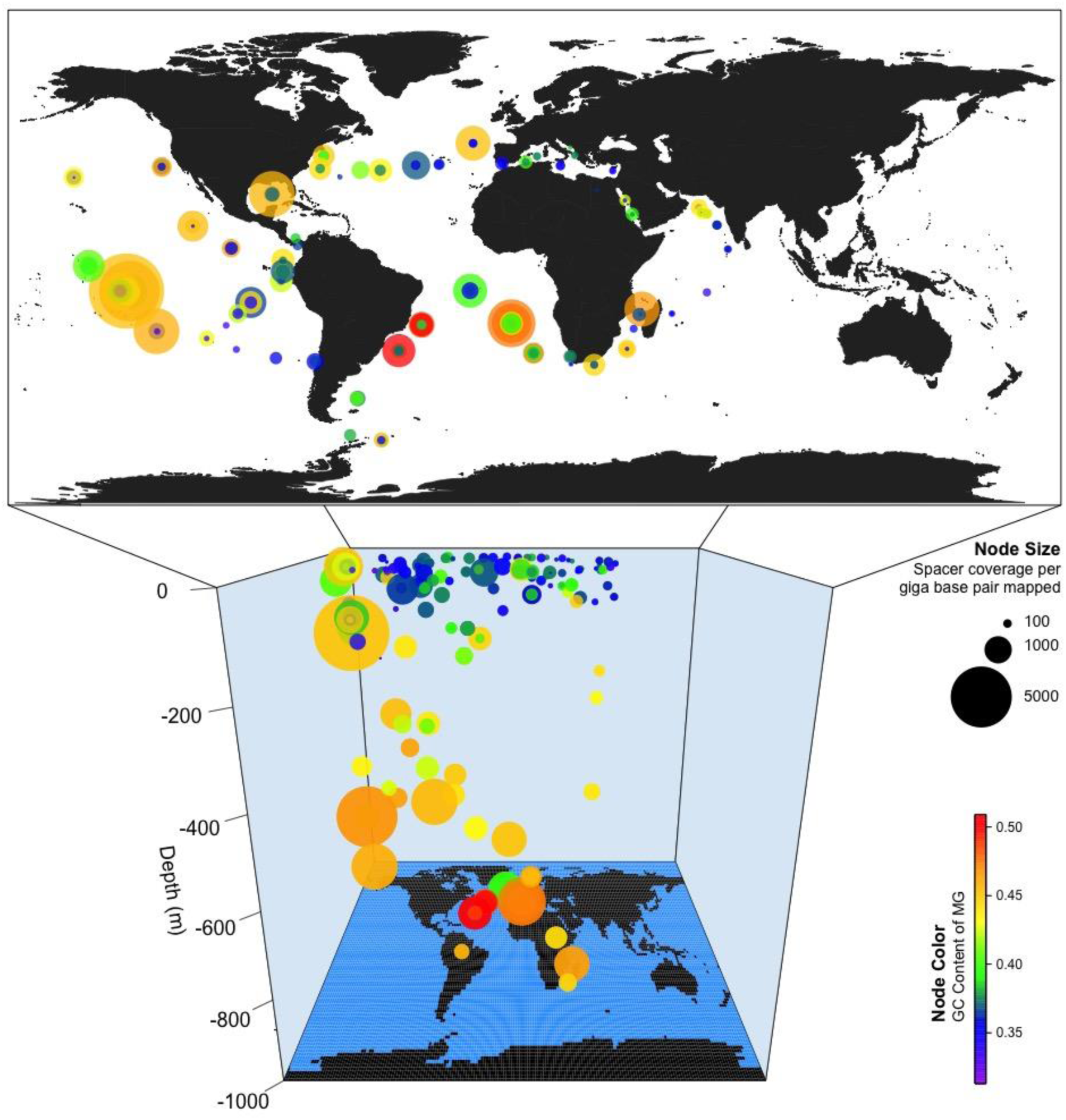
Normalized spacer abundance correlates with depth and GC content. Map of spacers found by *Tara* Oceans sites. Node size represents the normalized abundance of spacers at that *Tara* site (cumulative spacer coverage divided by mapped read gigabases for that sample), node color represents the mean GC content of contigs at that site.

### Linking CRISPR abundance to taxonomic composition of microbial communities

Observed CRISPR spacer abundances in the global oceans were analyzed with respect to the previously reported taxonomic composition of prokaryotic plankton communities within *Tara* Oceans metagenomes (22). Nearly 75% of archaeal 16S rDNA operational taxonomic units (OTUs) exhibited a positive correlation with NSA. Thus, as NSA increased, the abundance of archaeal OTUs was more likely to increase than decrease. In contrast only 50% of bacterial OTUs exhibited a positive correlation with NSA, meaning that as NSA increased the abundance of bacterial OTUs was equally likely to increase or decrease (p-value = 1.1e-14). Additionally, there was a positive correlation between NSA and Bray-Curtis dissimilarity, an index to assess microbial community similarity (Mantel r = 0.30, p-value = 0.01). Thus, the greater the compositional differences between prokaryotic plankton communities the greater the difference in their NSA values.

At varying depth zones, the SAR clades within the Alphaproteobacteria sub-phyla were consistently among the most negatively correlating OTUs with respect to NSA (Fig. 2). Interestingly, some taxa with OTUs that negatively correlated with NSA also had OTUs that positively correlated with NSA. In general, the percentage of OTUs with significant positive correlations to NSA increased with depth (Surface = 2.2%, deep chlorophyll maximum (DCM) = 6.6%, Mesopelagic = 7.5%), while the percentage of OTUs with negative correlations to NSA remained fairly steady with the exception of the DCM (Surface = 0.45%, DCM = 0.01%, Mesopelagic = 0.46%).

**Fig. 2.**
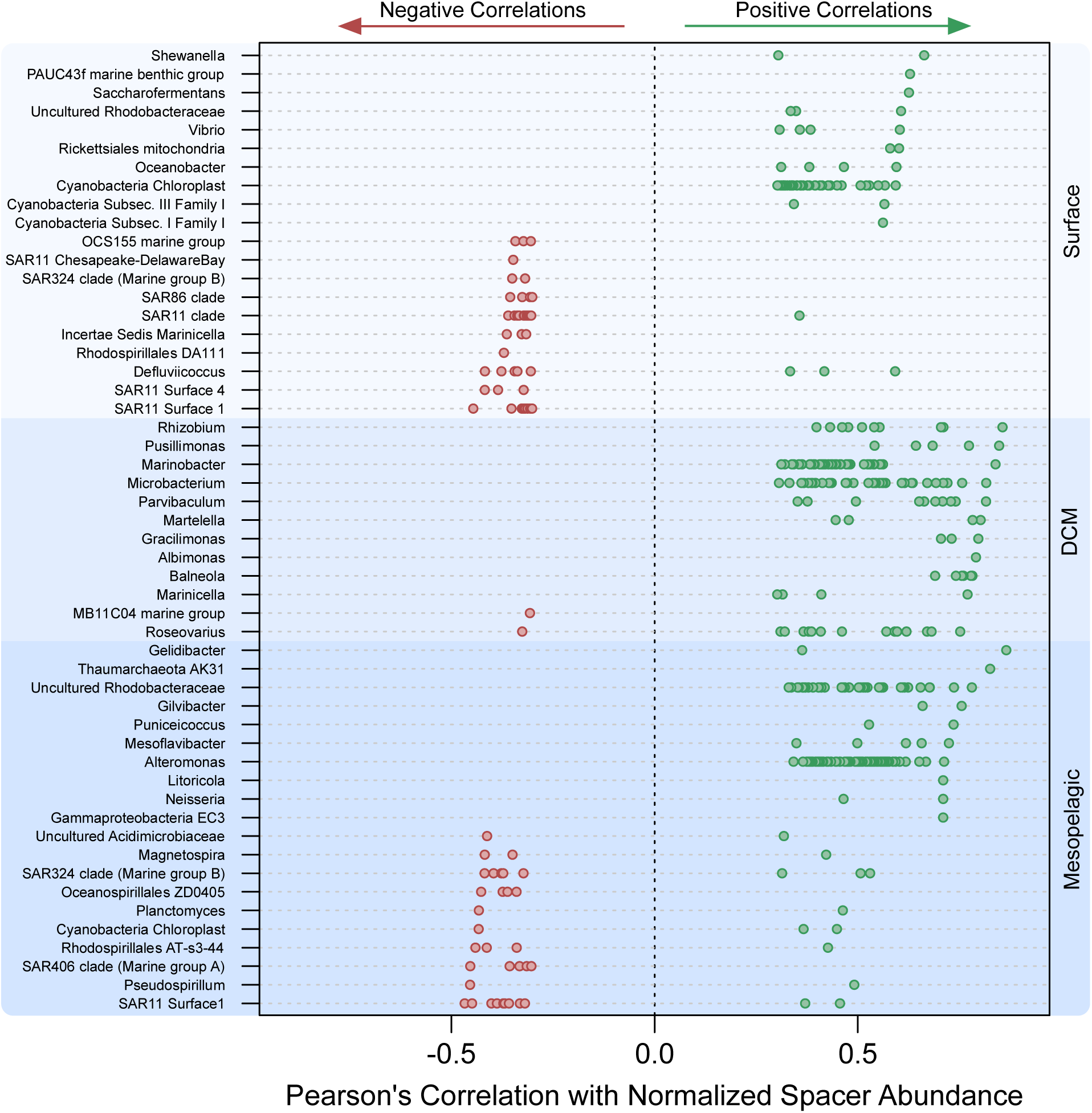
CRISPR abundance correlates with several taxonomic OTU’s, with stronger positive correlations in deeper ocean zones. The top 10 positively and negatively correlating OTUs with respect to normalized spacer abundance, broken down by oceanic depth zone. Some taxa have several significant OTUs.

### Some viral genes are more likely to become spacers

Matching a CRISPR spacer from a metagenome to a viral gene target (VGT) is challenging because: (*i*) the collection of known reference viral genomes poorly represents environmental viruses (especially aquatic viruses); (*ii*) viral genes mutate rapidly; and (*iii*) the short length of spacer sequences means that even alignments with a high percent identity match may have high BLASTn E values (Expect Values). To address these challenges, a large database of virome sequences comprising 206 aquatic viral metagenomes and totaling ca. eight giga base pairs (Gbp) of sequence data (65 *Tara* Oceans assembled viromes, 141 unassembled public viromes) was collected. All microbial spacers found in the GOS and *Tara* Oceans datasets were searched against the virome database with BLASTn (E value ≤ 1e-1, word size 7) to identify matches between spacers and candidate VGTs. Nucleotide open reading frames (ORFs) were predicted only for virome sequences with a spacer match, allowing for the detection of spacers that spanned two adjacent ORFs, which proved to be rare (3% of spacers).

A many-to-many relationship between CRISPR spacers and their candidate VGTs was observed – i.e. some spacers showed homology to multiple virome ORFs, and some virome ORFs showed hits from multiple spacers (Fig. 3). While the majority of spacers were homologous to only one virome ORF (nearly 1,500 spacers, 45%), there were a few spacers with homology to over 400 virome ORFs. These cosmopolitan spacers often targeted less complex regions of structural proteins such as short glutamic acid repeats within a portal protein.

**Fig. 3.**
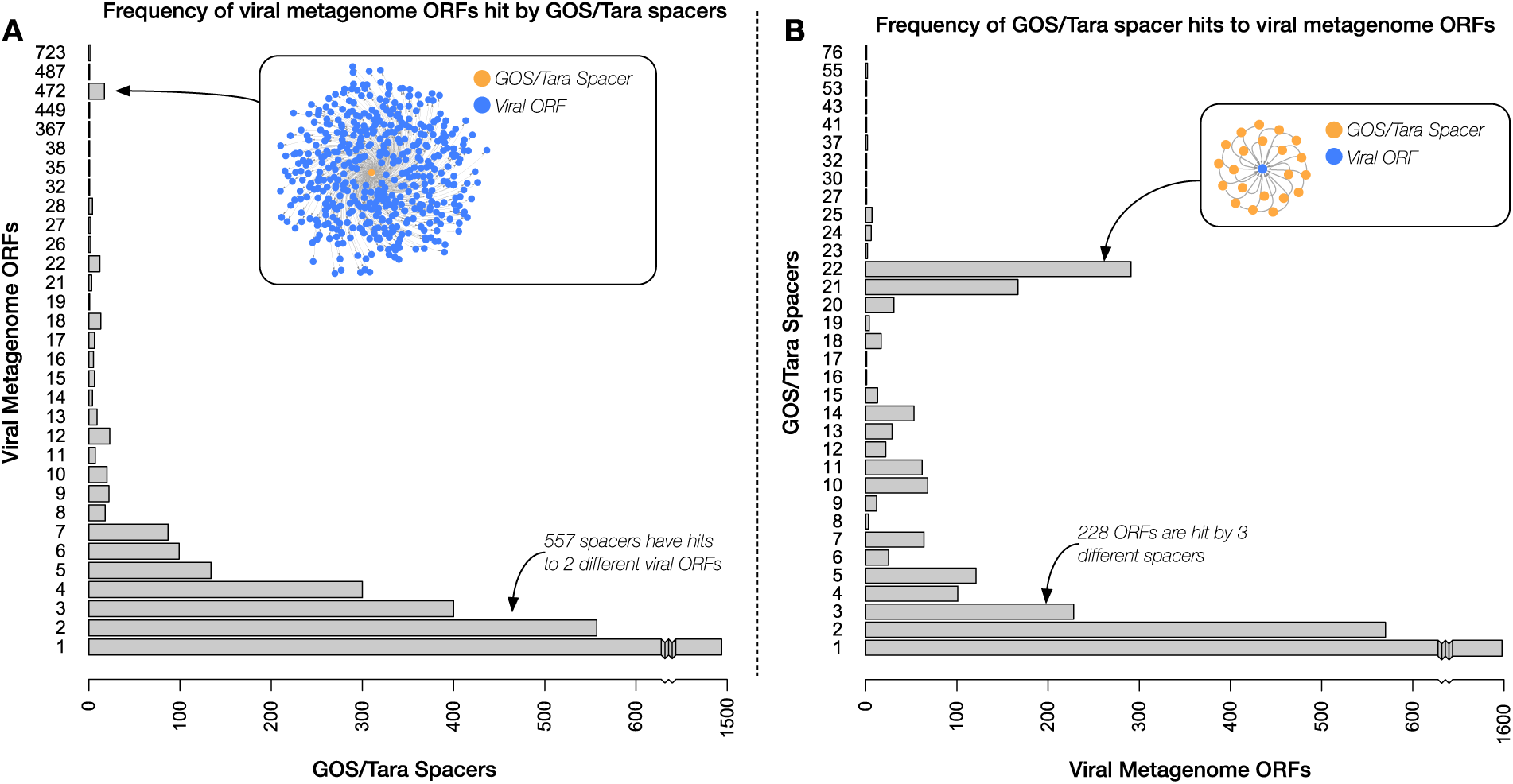
(A) CRISPR spacers aligned with viral gene targets in a many-to-many relationship. Frequency of viral metagenome ORFs hit by GOS and *Tara* Oceans spacers with inset network graph representing the 1 to 472 relationship. (B) Frequency of GOS and *Tara* Oceans spacer hits to viral metagenome ORFs with inset network graph representing the 22 to 1 relationship.

In total, nearly a quarter (24%) of the CASC-identified (run this time in conservative mode ensuring these were bona fide spacers) bacterioplankton spacers had a nucleotide BLAST alignment with a virome open reading frame. Nearly half of the translated viral ORFs (43%) had a match to a Phage SEED peptide (23), the majority of which had an informative annotation; i.e. were not simply labeled “Phage protein” (Fig. 4).

**Fig. 4.**
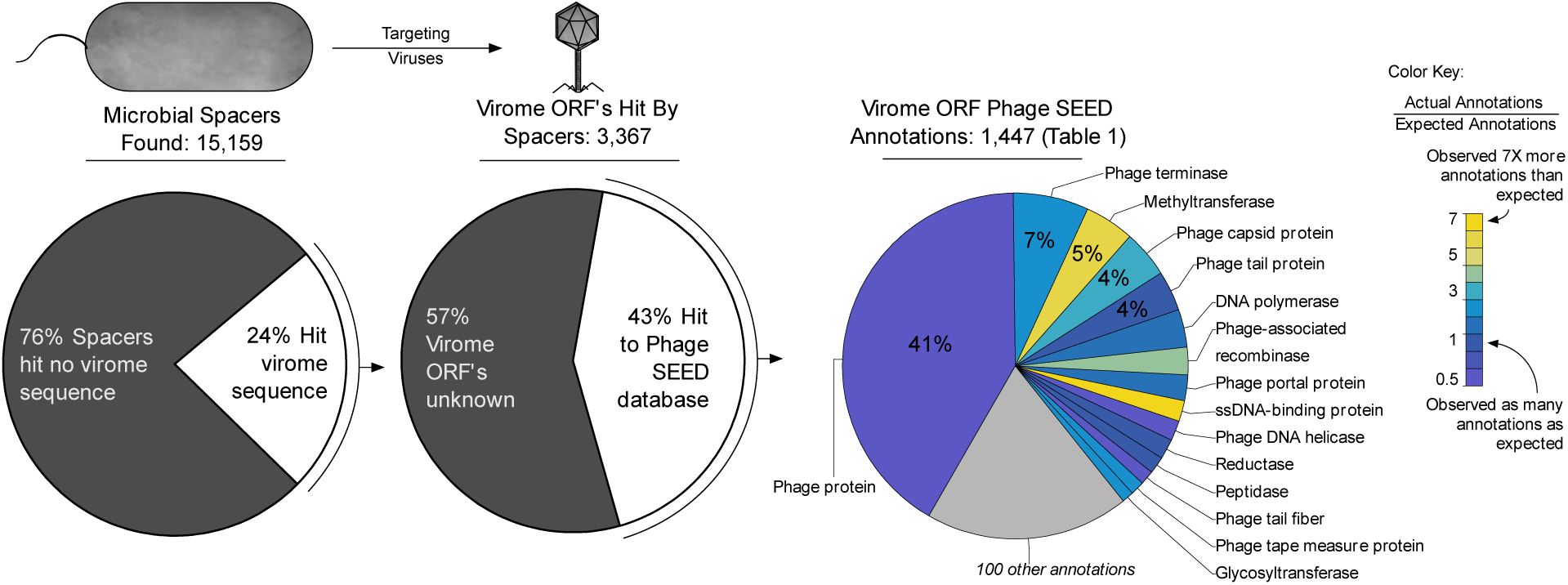
Microbial spacers preferentially target specific viral genes. Nearly one quarter of aquatic microbial spacers had a putative match to aquatic virome genes. The majority of these genes obtained informative annotations (i.e. not “Phage protein”). Most genes targeted by CRISPR spacers were annotated two-fold as often as expected, based on the expected frequencies of aquatic virome gene annotations. Two gene annotations that were seen less frequently than expected were DNA helicase and phage tail fiber.

All virome ORFs in the virome database were annotated using homology information to Phage SEED proteins, enabling quantification of the expected frequency of VGT annotations. In turn, annotation data was used to establish an expected frequency for each viral gene annotation within the collection global ocean viromes. Each of the top fifteen annotations assigned to VGTs were assigned more often than expected (Table 1, Additional file 5). There were two exceptions that were targeted less frequently than expected, genes encoding phage tail fiber (a set of structural proteins attached to the base of the tail, used in host recognition and attachment) and DNA helicase (a motor protein that separates double-stranded nucleic acid). Overall, the VGT ORFs had a higher rate of homology to Phage SEED peptides than would be expected indicating that VGTs of CRISPR defense are among the better-known subset of viral genes (expected 2,257 no-hits, observed 1,920).

**Table 1:** Fifteen most abundant virioplankton ORFs containing viral gene target sequences

| ORF Annotations | Actual Hits* | Expected Hits** | Fold (Act. / Exp.) |
| --- | --- | --- | --- |
| Phage protein | 598 | 699 | 0.9 |
| Phage terminase | 102 | 48 | 2.1 |
| Methyltransferase | 67 | 14 | 4.7 |
| Phage capsid protein | 64 | 25 | 2.5 |
| Phage tail protein | 54 | 47 | 1.2 |
| DNA polymerase | 51 | 32 | 1.6 |
| Phage-associated recombinase | 37 | 12 | 3.0 |
| Phage portal protein | 35 | 24 | 1.5 |
| ssDNA-binding protein | 28 | 4 | 7.1 |
| Phage DNA helicase | 27 | 37 | 0.7 |
| Reductase | 27 | 23 | 1.2 |
| Peptidase | 23 | 18 | 1.3 |
| Phage tail fiber | 19 | 28 | 0.7 |
| Phage tape measure protein | 19 | 9 | 2.0 |
| Glycotransferase | 16 | 8 | 1.9 |
| [104 Other annotations] | 272 | - | - |
| No hits | 1920 | 2257 | 0.8 |

\* Actual hits are the number of spacer hits
\*\* Expected hits were calculated based on the frequency of all aquatic viral ORFs being assigned a given annotation by Phage SEED

The *Tara* Oceans microbial shotgun metagenomes and viromes provided a rich set of spacer-to-virome ORF matches. However, instances of bacterioplankton spacers matching ORFs within a virome collected from the same water sample were rare. More frequently bacterioplankton spacers had matches to virome ORFs from viromes collected several thousand miles away (Fig. 5). This was the case for bacterioplankton metagenomes collected from surface and deep chlorophyll maximum water samples.

**Fig. 5.**
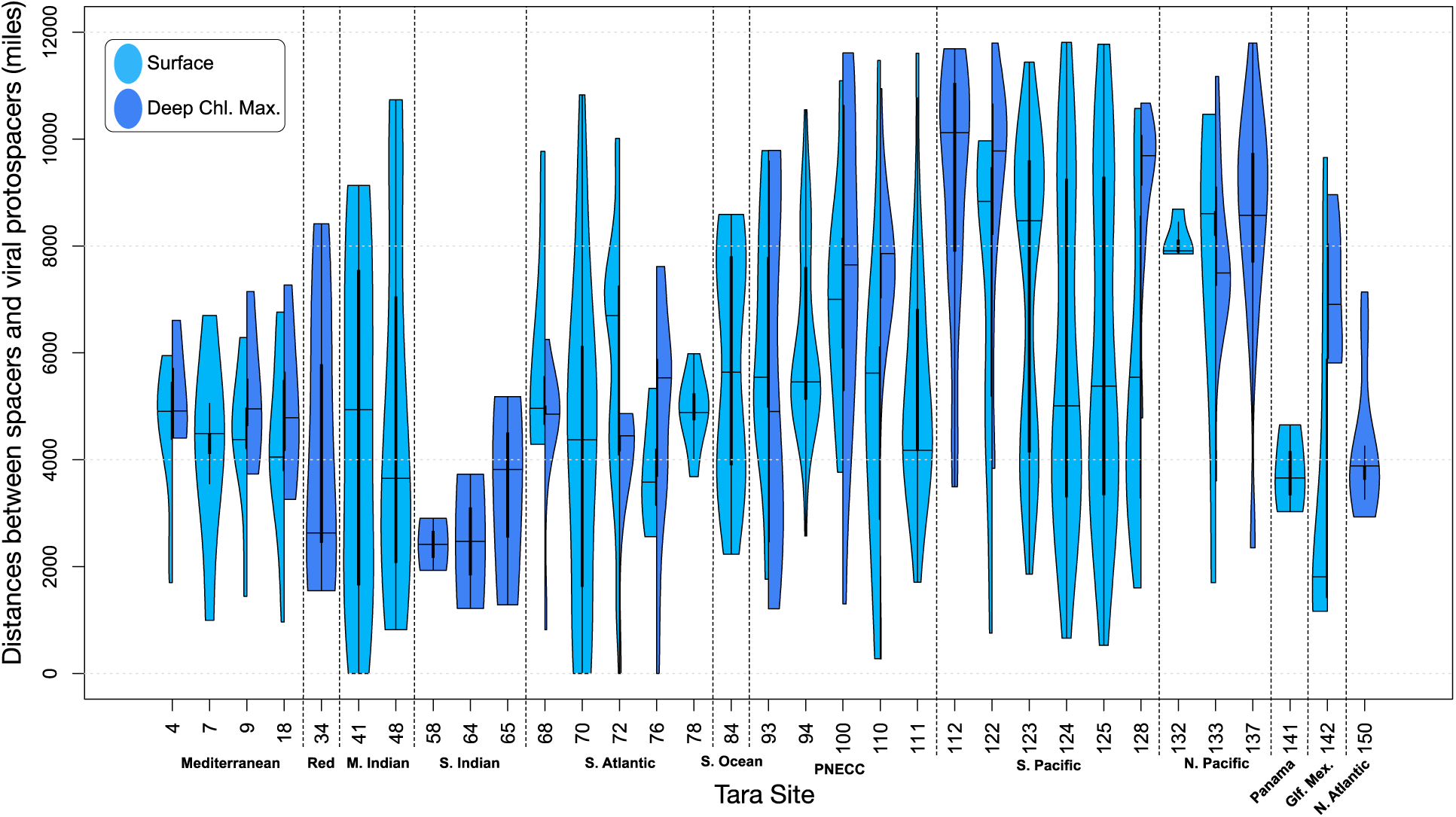
CRISPR spacers are more likely to match viral gene targets from distant viromes than paired viromes. Violin plots of the distances between *Tara* Oceans spacers and the viromes they aligned with (light blue, surface sample; dark blue, deep chlorophyll maximum sample). Line connections demonstrate sites with paired surface and DCM samples. Sites are broadly split by geographic location (Mediterranean, Mediterranean Sea; Red, Red Sea; M. Indian, Indian Monsoon Gyres; S. Indian, Indian S. Subtropical Gyre; S. Atlantic, S. Atlantic Gyre; S. Ocean, Southern Ocean; PNECC, Pacific North Equatorial Countercurrent; S. Pacific, South Pacific Ocean Gyre; N. Pacific, North Pacific Ocean Gyre; Panama, near Panama; Gf. Mex., Gulf of Mexico; N. Atlantic, North Atlantic Ocean Gyre).

### Viruses encoding CRISPR spacer arrays

Previous studies have shown that phages infecting marine bacteria can carry the genetic elements of the CRISPR/Cas system (24, 25). Over 2,000 CRISPR spacers were observed within the aquatic viromes. To determine if the virome spacers targeted a different subset of viral genes than the bacterioplankton spacers, the virome spacers were also assessed against the aquatic virome database, in the same way as the bacterioplankton metagenome spacers.

A greater frequency of virome spacers had a match to virome ORFs than that seen for bacterioplankton spacers (30% versus 24%). Additionally, more of these VGT ORFs of virome spacers could be annotated with Phage SEED than the bacterioplankton spacers (55% versus 43%) (Fig. 6, Additional file 6). Again, all of the ORFs in the virome database were annotated with Phage SEED to establish an expected frequency for each viral gene annotation in the global oceans. Among the informative annotations (annotations that were not simply “Phage protein”) methyltransferase was targeted 21 times more often than expected (expected ca. 5 annotations, observed 100) by viral spacers, whereas microbial spacers targeted methyltransferase only 4 times more often than expected. Indeed, methyltransferase was among several gene targets that are differentially targeted between microbial and virome spacers, including integrase and antitermination protein Q.

**Fig. 6.**
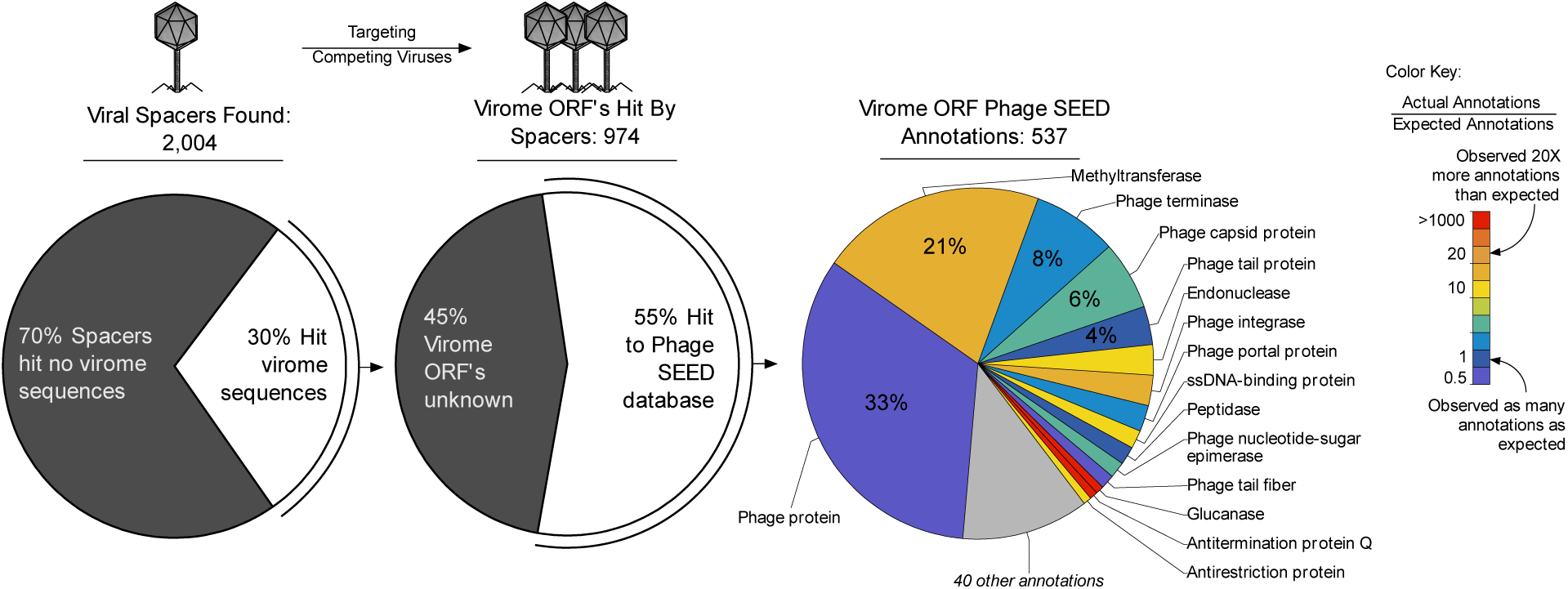
Over 2,000 CRISPR spacers were identified in the aquatic viral metagenomes and target methyltransferase more frequently than microbial spacers. Viral spacers are believed to assist the host in defending itself against competing viruses. Genes associated with temperate viruses (e.g. integrase, methyltransferase) are targeted more frequently by viral spacers than microbial spacers. Additionally, viral spacers targeted viral genes that were exceedingly rare in these aquatic dsDNA viromes, such as glucanase and antitermination protein Q, with many other genes being targeted >2X more often than expected. Again, phage tail fiber was targeted less frequently than expected.

## DISCUSSION

By and large, the focus of work investigating CRISPR as a microbial defense strategy has been to determine the biochemical mechanisms behind spacer acquisition and maintenance within bacterial (26) and archaeal (27) taxa. As a consequence these studies have been conducted in model organisms within experimental laboratory systems (28, 29), with some exceptions (30). Here we investigated the diversity and frequency of unknown CRISPR/Cas systems within the global ocean, an approach that broadly accounted for the influence of environmental selective pressures on the acquisition and maintenance of CRISPR spacers. These investigations revealed that particular subsets of virioplankton genes are highly targeted by the CRISPR defense system of bacterioplankton and that there is a many-to-many relationship of spacers to virioplankton genes.

Deeply sequenced shotgun bacterioplankton metagenomes enabled the search for novel CRISPR spacers across a wide geographic range of aquatic environments. Increasing sequence read lengths and yields from next generation sequencers have enabled modern assembly algorithms to better resolve the repeat-rich CRISPR locus (31) as seen through the high yield of CRISPR spacers in the *Tara* Oceans dataset. Testing indicated that the addition of quality control heuristics in CASC provided a more reliable set of CRISPR spacers than other CRISPR-finding algorithms.

With the rich set of CRISPR spacers mined directly from the environment it is possible to compare our findings to those obtained through mathematical theory and single-organism model systems. Normalized spacer abundance positively correlated with sample depth indicating that the CRISPR/Cas system is an important defense strategy for deep-sea bacterial and archaeal populations. The concentrations of hosts and viruses is known to decrease with depth in the ocean (32), thus, this observation agrees with previous work demonstrating that inducible immunity (i.e. CRISPR) is preferred in conditions where the concentrations of host and virus are low (33). Not only was NSA generally lower at the surface, where concentrations of hosts and viruses tend to be greater, there were also several surface water bacterioplankton taxa that exhibited strong negative correlations with NSA; chief among them were taxa within the abundant SAR11 clade (Pelagibacterales) (34). This may be further evidence of the limited effectiveness of CRISPR/Cas defense in competitive environments, as SAR11 members (notorious defense specialists) appear to favor other mechanisms of bacteriophage resistance (e.g. cryptic escape (35)) rather than CRISPR/Cas.

A protospacer is the 30-40 bp segment of a viral gene that is incorporated into a CRISPR array as a spacer. A motif, adjacent to the protospacer, called the PAM (protospacer adjacent motif) is essential to the spacer acquisition machinery (14) in Type I and II CRISPR/Cas systems. However considering the short and often degenerate nature of PAMs (e.g. 2 bp, 16-fold degenerate (36)), hundreds to thousands of potential PAM sites can exist within a viral genome. Thus, while the PAM plays a role in determining the site within a viral gene that becomes a protospacer it remains uncertain what, if anything, contributes to the retention of certain spacers within the array in a natural system. Given the commonality of PAMs within viral genomes, the most parsimonious explanation for the observed selection of particular VGTs within virioplankton metagenomes is positive selection pressure for effective viral defense. The CRISPR spacers observed within the bacterioplankton metagenomes were maintained because they were the most successful in minimizing the damaging impacts of viral infection and lysis on bacterioplankton populations. These data also provide interesting insights concerning those genes that are most critical to the processes of viral infection and lysis of bacterioplankton hosts.

In particular, these data show that there are conserved regions of potentially evolutionary constrained viral genes that are targeted more often than expected by CRISPR spacers from bacterioplankton populations. Genes encoding phage terminase (enzymes that initiate DNA packaging by cutting the DNA concatemer), methyltransferase (a family of enzymes that catalyze the transfer of a methyl group to DNA or RNA), recombinase (enzymes that catalyze exchanges of nucleic acid within a genome), and ssDNA-binding proteins (proteins that bind single-stranded DNA to prevent it from re-forming a double-stranded molecule) were among the most overtargeted genes within the virioplankton (Fig. 4). An inference from these observations is that these viral genes are under particularly stringent selection pressure which prevents the easy acquisition of point mutations that would ordinarily allow a viral gene target to evade spacer recognition, the critical first step in CRISPR defense. Thus, our analysis has pointed to particular gene functions that may have a heightened importance to successful replication of marine viral populations.

The observation of thousands of spacers within nearly 20% of the viromes surveyed (38 of 206) indicated a high prevalence of CRISPR-carrying viruses. The impact of CRISPR-carrying viral populations in natural microbial communities may be greater than expected. The frequent observation of virome spacers supports the recent finding that cyanophages have been shown to carry CRISPR arrays and perhaps transfer the arrays between related cyanobacteria to offer infection resistance from competing phage (25). An enrichment in viral spacers targeting methyltransferase and integrase genes may indicate that viral CRISPR arrays aid the host in targeting competing temperate phage.

Interestingly, CRISPR spacers from bacterioplankton metagenomes targeted certain genes less frequently than expected such as phage tail fiber genes. The relatively simple structure of phage tail fiber protein would indicate a less stringent selective pressure at the coding level, implying a greater opportunity for tail fiber gene diversity. Indeed, phage tail fiber genes have been shown to not only be hypervariable, but also undergo targeted hypervariation by retroelements in order to expand viral host range (38, 39). Additionally, viral ORFs targeted by CRISPR spacers were less likely to have an unassigned function than expected (actual unassigned functions = 598, expected = 699) indicating CRISPR-targeted viral genes are more likely to have a known functional role as opposed to non-targeted genes (Fig. 4 and Table 1). Nevertheless, nearly half (41%) of these CRISPR-targeted viral genes were unknown and would be considered viral genetic “dark matter” (40). This subset of CRISPR-targeted but unknown viral “dark matter” genes likely play an important role in infection and lysis processes.

Spacers matched virome ORFs in a many-to-many relationship, indicating that some spacers were capable of targeting several different virome ORFs and several virome ORFs were targeted by multiple spacers. In the latter case, these viral genes appear to be highly targeted by the CRISPR/Cas system (Fig. 3). Instances of virome ORFs being targeted by multiple spacers suggests that these ORFs are under especially stringent selection pressure and are thus less likely to evade CRISPR interference through single nucleotide point mutations. The over-targeting of these ORFs also indicates that they are critical to viral replication and are thus more effective targets for bacterioplankton CRISPR immunity.

Interestingly, less than 1% of spacers from *Tara* Oceans microbial metagenomes matched virome ORFs from the same site (Fig. 5). One potential explanation for this observation is that spacers found in a given bacterioplankton metagenome have successfully minimized the replication of targeted viral populations to a level below detection within a virome library. This observation is consistent with previous studies of Archaeal-dominated systems (41, 42) and emphasizes a potential challenge of using CRISPR to link viruses with their hosts within a single environmental sample. The analysis of paired microbial/viral metagenomes over time may provide interesting perspectives, as it could be possible to observe spacers targeting viruses from past samples.

This study analyzed a large collection of CRISPR spacers from microbial populations throughout the global oceans and has provided evidence that particular viral genes are preferentially targeted by the CRISPR/Cas system. The identification of certain viral gene classes that are more likely to become CRISPR spacers indicates that these genes represent a genetic vulnerability for viral populations and that these genes are potentially under strict selective pressure for successful viral infection and lysis. CRISPR spacers sequenced from the environment have shown to be useful in linking microbial hosts to their viruses (43). Our findings also indicate that spacer sequences can identify those viral genes that represent the points of greatest genetic vulnerability for natural viral populations. In this way, CRISPR/Cas may be thought of as a living “evolutionary algorithm” (a field of artificial intelligence, which mimics natural selection to solve complex problems) to agnostically identify viral genes that are most vulnerable. These genes may then be further explored for uses in biotechnology (e.g. preventing phage infections in processes relying on bacterial fermentation) or analysis of phage diversity (as they are likely conserved).

## METHODS

### CASC Pipeline

The CASC pipeline can be broadly divided into two parts (Fig. S1): (A) preliminary search for putative CRISPR spacers and (B) validation of putative CRISPR arrays by Cas protein homology, CRISPR repeat homology, and the statistical characteristics of spacer sizes. The preliminary search for CRISPR arrays employs a modified version of the CRT (16). Modifications included a reformatting of the search output, improved handling of multi-FASTA files, and the ability to utilize multiple CPUs to lessen computational run time. These modifications improved the ability of CRT to analyze large metagenomic datasets. Putative CRISPR arrays are then validated and deemed “bona fide” CRISPRs if any of the following conditions are met: (*i*) the sequence containing the candidate CRISPR array has a BLASTx match (E value ≤ 1e-12) to a known UniRef 100 Cas protein cluster (44), (*ii*) the candidate CRISPR repeat had a BLASTn match (E value ≤ 1e-5, word size 4) to a known CRISPR repeat from the CRISPRdb reference database (7), or (*iii*) the standard deviation of spacer length within the candidate CRISPR array was less than or equal to two base pairs. CASC offers a “conservative” and a “liberal” CRISPR validation mode. In conservative mode, conditions (*i*) or (*ii*) must be met, while in liberal mode conditions (*i*), (*ii*), or (*iii*) may be met. CASC is available on GitHub (https://github.com/dnasko/CASC).

### Simulated Metagenome Construction

Two shotgun sequence simulations were generated using Grinder (ver. 0.5.0) (45) for the purpose of validating CASC and assessing performance. Ten complete bacterial genomes were selected for the simulated metagenomes (Table S1), five of which contained CRISPR arrays. The first simulation generated 60 million paired-end 150 base pair Illumina reads (read_dist=150 normal 0; insert_dist=300; mutation_dist=poly4) and the second simulation generated 1 million 454 pyrosequencing reads (read_dist=450 normal 50; mutation_dist=poly4).

The Illumina simulated read pairs were assembled using the St. Petersburg genome assembler (SPAdes) version 3.5.0 using all default settings (19) with the exception of bypassing the pre-assembly read error correction process. The 454 simulated reads were not assembled and CRISPRs were predicted directly from the reads.

### Performance Validation

The known CRISPR array positions in five of the ten genomes were used to assess the performance (i.e. sensitivity and precision) of several CRISPR identification algorithms. Alignment of the Illumina assembled contigs against the reference genomes identified the position of each CRISPR locus on the contigs and indicated that all spacers were successfully assembled. The alignment-generated CRISPR positions on the contigs were then used as the known CRISPR array positions. CRISPR array positions within the 454 reads were determined using the genome coordinates provided by Grinder.

Several algorithms, including CASC version 2.5 and the default settings of metaCRT (a version of the CRT modified by Rho and colleagues) (46), PILER-CR (ver. 1.06) (20), and CRISPR Finder (21), were used to predict CRISPR arrays from the Illumina assembled contigs and 454 reads (Table S2 and Table S3). Predicted spacers from each program were clustered with the set of known spacers using CD-HIT-EST (ver. 4.6) (47). Those spacers clustering at 100% identity with a known spacer were counted as a true positive.

To better measure the abundance of spacers in the simulated Illumina metagenome a recruitment of the simulated Illumina reads to assembled SPAdes contigs was performed using Bowtie2 (ver. 2.1.0) (48). Coverage of each spacer was calculated using SAMtools (ver. 1.2-2-gf8a6274) (49) and used to estimate the number of spacer copies present in the simulated Illumina metagenome.

### Spacer predictions in GOS and *Tara* Oceans microbial metagenomes

The Global Ocean Sampling (GOS) and *Tara* Oceans expeditions sampled and sequenced microbial DNA from across the world’s oceans (17, 18). The GOS dataset was ideally suited for CRISPR prediction as the long read technology used for sequencing these libraries was capable of encoding intact CRISPR arrays (50), and this dataset has been used in previous studies of CRISPR prediction from metagenomic data (51, 52). GOS sequences were downloaded from iMicrobe (imicrobe.us) and included the GOS I expedition, GOS Baltic Sea, and GOS Banyoles (Additional file 1). CRISPR spacers were predicted from 157 GOS sequence libraries totaling ca. 39 million reads and containing ca. 21 Gbp of genomic DNA from microorganisms typically between 0.1 and 0.8 μm in size (note that filter sizes ranged from 0.002 to 20 µm based on sample site) with CRISPR calling in ‘liberal’ mode.

The *Tara* Oceans expedition was a global-scale oceanic study that sampled and sequenced metagenomes from 67 sites (53). In addition to sampling nearly every site at varying depths, several sites were processed with multiple filter sizes (ranging from 0.2 to 3.0 µm), including 54 sites with paired microbial and viral fractions, making the *Tara* Oceans dataset ideal for linking bacterial spacers with their viral gene targets in the viromes. *Tara* Oceans metagenomes were predominantly sequenced using Illumina HiSeq (100 bp, paired-end reads). Because Illumina reads are too short for accurate searches of spacer arrays, assembled contigs were used instead (ca. 58 million contigs totaling 62 Gbp). *Tara* Oceans assembled contigs were obtained from the European Nucleotide Archive (http://www.ebi.ac.uk/ena/about/tara-oceans-assemblies).

In addition to counting the number of spacers found within each *Tara* contig, it was necessary to calculate the abundance of each spacer by recruitment of the original library of unassembled Illumina reads to *Tara* contigs. The reads corresponding to each assembly were downloaded from NCBI’s Sequence Read Archive and recruited to their assembled contigs using Bowtie2 (very sensitive local setting). Read coverage of each spacer was calculated using SAMtools and used as a proxy for the number of copies of each spacer.

To measure how novel these spacers were, the GOS and *Tara* Oceans spacers were clustered with known spacers from the CRISPRDB at 98% identity using CD-HIT-EST (7, 47).

### Microbial Community Profiles with Respect to CRISPR Abundance

The *Tara* Oceans observed OTUs “16S OTU Table” from Sunagawa et al. (22) was downloaded from http://ocean-microbiome.embl.de/companion.html and imported into QIIME (54). OTUs occurring ≤ 2 times were filtered out and 100 jackknife subsamples were created with 35,461 observations (90% of the smallest sample) in each. The community similarity test was performed with beta_diversity.py using Bray-Curtis. Per-OTU correlations were calculated for each depth zone after splitting the BIOM file accordingly and using observation_metadata_correlation.py. Only correlations with Pearson’s r ≥ 0.3 or ≤ −0.3 with p-value ≤ 0.05 were considered significant.

### Identification of GOS and *Tara* Oceans Spacer Targets

Putative CRISPR spacers from the GOS and *Tara* Oceans microbial metagenomes were searched against *Tara* Oceans viromes (Additional file 3) and a subset of publicly available aquatic viromes (Additional file 4) available on the Viral Informatics Resource for Metagenome Exploration (VIROME, virome.dbi.udel.edu) (55) to identify candidate viral gene targets. Only spacers found with CASC in conservative mode were used for this analysis to reduce the likelihood of identifying spurious spacers.

Sequence alignment cut-offs used in previous studies comparing microbial spacers to virome genes have varied, both in stringency and cut-off metric, depending on the aim of the study. When identifying host-phage interactions by linking specific viral population(s) to CRISPR spacers/loci, more stringent cut-offs are applied, such as requiring a 100% nucleotide identity alignment of ≥ 20 bp (11), or an alignment with no more than one mismatch (56). Exploratory studies trying to link what, if any, similarities exist between microbial spacers and virome genes have used more relaxed cut-offs, such as E value ≤ 1e-3 (10), or alignments containing up to 15 mismatches (57).

As the objective of this study was to determine if particular viral genes were more likely to be targeted by the CRISPR system of marine bacterioplankton the latter, more exploratory approach was used. Spacer sequences are highly diverse and hyper variable, even between closely related species (58), making it challenging to identify candidate viral gene targets at the nucleotide level. Thus, when searching for potential viral gene targets in viromes some mismatches and gaps in the nucleotide alignment were permitted using BLASTn (ver. 2.2.30+, E value ≤ 1e-1, word size 7). This resulted in 51% of high-scoring segment pairs (HSPs) with no mismatches and 89% of HSPs with no gap openings (Fig. S3).

In this analysis some spacers matched CRISPR arrays within several viromes. To limit these spurious matches, CASC (liberal mode) was used to identify putative spacer arrays within the viromes. Subsequently, sequences containing an array were removed from the aquatic virome database prior to the analysis to identify viral gene targets.

Spacer sequences were searched against the virome database with BLASTn. Virome sequences that aligned with spacers were then culled into a separate FASTA file and open reading frames (ORFs) were predicted using MetaGene (59). ORFs were predicted after the spacer search to detect any spacers that may have spanned virome ORFs (a rare occurrence). Virome ORFs with a match to a spacer were translated and searched against Phage SEED (version 01-May-2016) (http://www.phantome.org) using BLASTp (ver. 2.2.30+, E value ≤ 1e-3). Each ORF was annotated using the best cumulative bit score, which is described in the next section.

Finally, great-circle distances between microbial metagenome spacers and VGTs within viromes were calculated in R (60) using the geosphere package (61). Distance distributions were rendered in violin plots using the R package vioplot.

### Annotating virome ORFs and calculating expectation

Virome ORFs with a match to a spacer were translated and searched against Phage SEED (version 01-May-2016) (http://www.phantome.org) using BLASTp (ver. 2.2.30+, E value ≤ 1e-3). A virome ORF was annotated to be the gene function producing the highest cumulative bit score. For example, if “ORF_1” hit ten Phage SEED genes, eight of which were hits to phage protein and the total bit score of these alignments was 50, while the two remaining hits were to terminases with a total bit score of 100, then “ORF_1” would be assigned to terminase. ORF annotation counts were generated for the virome ORFs matching microbial (Additional File 5) and virome spacers (Additional File 6).

To put these counts in come context, all aquatic virome ORFs were run through the same Phage SEED-based annotation pipeline. Counts for all virome ORFs were tabulated and the frequency of occurrence for each gene type was calculated. The expected number of genes to have matches to CRISPR spacers was calculated by multiplying the total number of genes matching spacers by the frequency of that gene being annotated in all aquatic viromes.

## Data Availability

Scripts used in this analysis are available on GitHub (github.com/dnasko/CASC) under the GNU General Purpose License.

Six datasets were used in this analysis. The first two were simulated metagenomic datasets and are available at Zenodo (http://doi.org/10.5281/zenodo.1650429). The second two datasets were shotgun metagenomic reads from the Global Ocean Survey (GOS) and *Tara* Oceans survey. GOS sequences were downloaded from iMicrobe (imicrobe.us) and included the GOS I expedition, GOS Baltic Sea, and GOS Banyoles (Additional file 1). *Tara* Oceans assembled contigs were obtained from the European Nucleotide Archive (http://www.ebi.ac.uk/ena/about/tara-oceans-assemblies). The fifth dataset was a subset of publicly available aquatic viromes (Additional file 4) available on the Viral Informatics Resource for Metagenome Exploration (VIROME, virome.dbi.udel.edu). Finally, the *Tara* Oceans observed OTUs “16S OTU Table” from Sunagawa et al. (22) was downloaded from http://ocean-microbiome.embl.de/companion.html.

## Acknowledgements

This work was supported through grants to KEW and SWP from the National Science Foundation (OCE-1148118, OIA-1736030 and DBI-1356374), the National Institutes for Health (5R21AI109555-02) and the Gordon and Betty Moore Foundation (grant number 2732). Support from the University of Delaware Center for Bioinformatics and Computational Biology Core Facility and use of the BIOMIX compute cluster was made possible through funding from Delaware INBRE (NIGMS GM103446) and the Delaware Biotechnology Institute.

## Authors’ Contributions

D.J.N, S.W.P., and K.E.W designed research; D.J.N. and R.M.M performed the research; D.J.N. and J.D.B. wrote and modified software and D.J.N. and K.E.W. wrote the paper. D.J.N., B.D.F., S.W.P., and K.E.W. revised the paper.

## SUPPLEMENT

**Figure S1**: The CASC Workflow. A) Preliminary search for CRISPR arrays and identification of putative spacer arrays. B) Validation of putative spacers.

**Figure S2**: Nucleotide position histogram of CRISPR repeats from (A) CRISPR repeats deemed “bona fide” by CASC, (B) all CRISPR repeats from CRISPR DB, and (C) CRISPR repeats deemed Non-”bona fide” by CASC.

**Figure S3:** Alignments between spacers and viral ORFs were typically strong. (A) Nearly 95% of HSPs had 3 or fewer mismatches in alignments of spacers to viral ORFs. (B) Nearly 98% of HSPs had 1 or no gaps open in alignments between spaces and viral ORFs.

**Additional File 1**: CRISPR spacers found in GOS datasets

**Additional File 2**: CRISPR spacers found in the Tara Oceans microbial metagenomes

**Additional File 3**: Summary of Tara Oceans viromes

**Additional File 4**: Summary of aquatic viromes collected from VIROME (virome.dbi.udel.edu)

**Additional File 5**: Actual vs. expected number of annotations for candidate microbial-viral protospacers

**Additional File 6**: Actual vs. expected number of annotations for candidate viral-viral protospacers

**Table S1.** Bacterial genome sequences used in the construction of the mock metagenomes.

| Organism | %<br>GC | Genome<br>Size (Mbp) | Spacers | Arrays |
| --- | --- | --- | --- | --- |
| *Escherichia coli* str. K-12 substr. MG1655 | 51 | 4.6 | 18 | 2 |
| *Streptococcus salivarius* JIM8777 | 40 | 2.2 | 32 | 1 |
| *Neisseria meningitidis* 8013 | 51 | 2.3 | 25 | 1 |
| *Yersinia pestis* A1122 | 48 | 4.5 | 16 | 3 |
| *Chlorobium tepidum* TLS | 57 | 2.1 | 62 | 2 |
| *Chlamydia trachomatis* F/SW5 | 41 | 1.0 | - | - |
| *Ruegeria pomeroyi* DSS-3 | 64 | 4.1 | - | - |
| *Bacillus thuringiensis* HD-789 | 35 | 5.5 | - | - |
| *Bordetella pertussis* CS | 68 | 4.1 | - | - |
| *Acetobacter pasteurianus* IFO 3283-01 | 53 | 2.9 | - | - |

**Table S2.** CRISPR finding tool performance Spacers found in the artificial 454 pyrosequencing metagenome using available CRISPR discovery tools.

| Program | Spacers<br>in Dataset | Spacers<br>Detected | True<br>Positives | False<br>Positives | False<br>Negatives | Sensitivity | Precision |
| --- | --- | --- | --- | --- | --- | --- | --- |
| CASC - Conservative | 1930 | 981 | 802 | 179 | 1128 | 0.42 | 0.82 |
| CASC - Liberal | 1930 | 1623 | 1108 | 515 | 822 | 0.57 | 0.68 |
| CRISPR Finder | 1930 | - | - | - | - | - | - |
| metaCRT | 1930 | 2631 | 1225 | 1406 | 705 | 0.63 | 0.47 |
| PILER-CR | 1930 | 1483 | 1070 | 413 | 860 | 0.55 | 0.72 |

**Table S3.** Spacers found in the artificial Illumina metagenome using available CRISPR discovery tools.

| Program | Spacers in<br>Dataset | Spacers<br>Detected | Spacer<br>Coverage<br>in Dataset | Spacer<br>Coverage<br>Detected | True<br>Positives | False<br>Positives | False<br>Negatives | Sensitivity | Precision |
| --- | --- | --- | --- | --- | --- | --- | --- | --- | --- |
| CASC - Conservative | 153 | 153 | 42,349 | 41,095 | 153 | 0 | 0 | 1.00 | 1.00 |
| CASC - Liberal | 153 | 153 | 42,349 | 41,095 | 153 | 0 | 0 | 1.00 | 1.00 |
| CRISPR Finder | 153 | 216 | 42,349 | 62,418 | 153 | 63 | 0 | 1.00 | 0.71 |
| metaCRT | 153 | 365 | 42,349 | 96,503 | 153 | 212 | 0 | 1.00 | 0.42 |
| PILER-CR | 153 | 146 | 42,349 | 39,222 | 138 | 8 | 15 | 0.90 | 0.95 |

